# Tuning Spatial Distributions of Selection Pressure to Suppress Emergence of Resistance

**DOI:** 10.1101/2024.10.23.619847

**Authors:** Thomas Tunstall, Philip G. Madgwick, Ricardo Kanitz, Wolfram Möbius

**Affiliations:** Living Systems Institute, Faculty of Health and Life Sciences, University of Exeter, Exeter, United Kingdom; Physics and Astronomy, Faculty of Environment, Science and Economy, University of Exeter, Exeter, United Kingdom; Syngenta, Jealott’s Hill International Research Centre, Bracknell, United Kingdom; Syngenta Crop Protection, Basel, Switzerland

## Abstract

Control measures such as insecticides or antimicrobials are used to contain biological agents such as pathogenic bacteria and vectors of human and plant diseases, respectively. Following control measure application, a resistant subpopulation may eventually rise to such frequency that the control measure will be rendered ineffective: The timescale over which this occurs is the ‘effective lifetime’ of the control measure. Prolonging this timescale relaxes urgency at which novel control measure needs to be developed. Spatial heterogeneity in control measure application can influence the rate at which resistance to the control measure evolves; in the agricultural context, this fact is exploited by distributing insecticides in mosaics across cropping regions in order to slow the rate of resistance evolution.

Contemporary and historical modeling practices, which aim to inform agricultural practices, often employ assumptions which squeeze out the impact of the spatio-temporal heterogeneity endemic to nature. In this paper, we present a minimal model of continuous dispersal and spatio-temporal heterogeneity in selection pressure distribution which exhibits a novel dynamic: The spatial distribution of selection pressure may be tuned in order to minimize the initial rate at which resistance evolves, thus increasing the effective lifetime of a pesticide.

**Author summary:** There are many contexts in which humans apply control measures to biological agents: pesticides are applied in fields to kill the pests that damage crops, mosquito nests are distributed to prevent the spread of malaria, and cancer drugs are applied to kill off tumors in humans. These control measures are examples of ‘selection pressures’, which select against strains which are susceptible to them. If a mutation occurs which confers resistance, the control agent will select for the resistant strain, thus reducing the efficacy of the control agent as the frequency of resistance increases in the population. This necessitates more of the control agent to be applied, or for a novel control agent to be developed. The former may have unintended consequences on the local environment, and the latter is expensive and time-intensive. It is preferable to carefully tune *how* the control agent is applied - perhaps instead of one massive compact region of control measure application, it is better to apply the control measure over multiple smaller regions? Here, we use a toy model of motile organisms to demonstrate that an optimal distribution exists for a variety of scenarios.

## Introduction

As an agricultural pest, arthropods in general, but primarily insects, cause approximately 20% of global crop losses [SKA17]. Insects also propagate diseases: *Anopheles* mosquitoes transmit malaria, resulting in 247 million cases of malaria and 619, 000 deaths in 2021 alone [WHO+22]. Insecticides are employed to control these pest populations. However, the application of such control agent results in the spread of resistance in the population [Met89]. This in turn acts to reduce the effective useful lifetime of the insecticide, thereby begging the question of how this lifetime can be extended by reducing the rate at which insecticide resistance evolves.

One solution is to apply insecticides not uniformly, but in a ‘mosaic’ fashion. The idea behind this approach is that regions free from insecticide application act as regions of ‘refuge’ from which the susceptible population can maintain high frequency and thus dilute the resistant population, thus reducing the emergence of resistant pests. In the literature, this approach is frequently compared against alternative possibilities, such as temporal heterogeneity in insecticide application, the application of multiple insecticides at the same time, or any combination of these [Rim+18; Con+13].

Assessment of optimal application procedures is typically supported by modeling, which is often self-described as simplistic or heuristic [SH18; MK22] in order to reduce computational complexity while still capturing important features of the real system. Many models make assumptions that suggest environments are homogeneous in both time and space: For example, when it is assumed that there is homogeneity in insecticide application over the entire landscape for the entire generation [Man85; Com86]. Even when considering mosaic models, fine-grained features of spatio-temporal dispersal are repeatedly excluded from modeling approaches: For example, having dispersal only occur in one event per generation [Cur85], or permitting dispersal to occur arbitrarily far over space [GT77; Com77; Man89]. It is under such simplifying assumptions that it has been determined that standalone mosaic application is one of the less successful methods to inhibit resistance emergence.

Work that does account for the continuity in space and time in spatio-temporally heterogeneous environments demonstrate that spatially structured regions of insecticide application and refuge can be tuned to minimize resistance emergence [Mui75] and overall pest density [LR98] better than a blanket application of insecticide. Based on the assumption that there is a cost to resistance, this previous work focused on minimizing the overall emergence of resistance in the absence of de novo mutations. While there are many documented cases in which there is a significant fitness cost to resistance [KG12], it has also been noted that some resistance costs are small or negligible [Bas+17]. Therefore, the case of negligible cost to resistance is an important limiting case to consider, and is fertile ground for a determination of optimal mosaic structure to minimize the initial rate at which resistance arises. In this work, we demonstrate with a minimal model of insecticide application (which considers fine-grained spatial features and no fitness cost to resistance) that insecticide application can be spatially optimized for reduced rate of emergence of resistance.

While motivated by insecticide application, the model is general and does not incorporate specifics of pests or insecticides. It can thus be more broadly described as a general minimal model for evolution occurs in any scenario in which motile agents move in a spatio-temporally heterogeneous landscape of control measure. Hence, the results of this work are presented generally in terms of tuning a ‘selection region’ distribution: A selection region may correspond to a region with pesticide application in the agricultural context, or a high prevalence of antibiotics or mosquito nets in an epidemiological context [Con+13].

As we determine how to tune the spatial distribution of selection region, we explore three distinct geometries in 1D. First, in the limiting case where regions of control agent application are separated infinitely far from each other, we show that the size of ‘isolated’ regions of control agent can be optimized to minimize the rate of resistance evolution, using computational, numerical, and analytical arguments (Fig. 1ci). Second, as another limiting case where many regions of control agent application are distributed close to each other, we find an optimal arrangement for an infinitely ‘periodic’ sequence of regions of control agent and refuge (Fig. 1cii). Finally, we investigate cases where finitely many or ‘multiple’ regions of control measure are evenly distributed over finite space (Fig. 1ciii). The results of this more complex scenario can be well-approximated by the first two cases in certain regimes, demonstrating that the results obtained in this work are useful heuristics for studying more realistic distributions of control measures.

**Fig 1.**
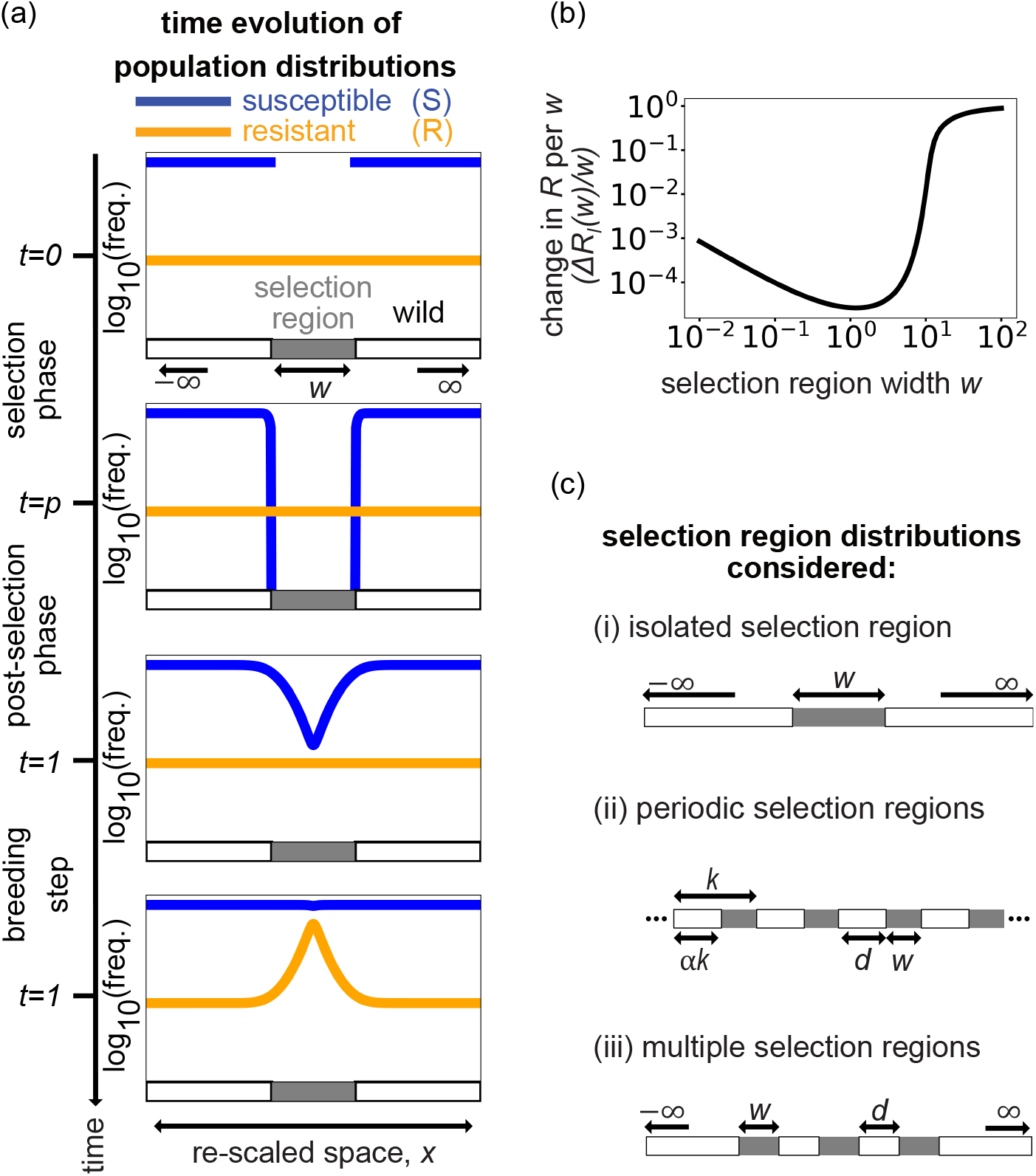
Overview and visualization of model and optimum size of selection region. **a)** Visualization of population densities *u*_*S*_(*x, t*) (blue) and *u*_*R*_(*x, t*) (orange) for susceptible and resistant individuals, respectively, over the course of the generation, i.e., at start and end of selection phase as well as before and after breeding for *p* = 0.1 and *ϕ* = 10^−5^. **b)** Change in the number of resistant individuals Δ*R* over one generation normalized by selection region size *w* for an isolated selection region corresponding to panel (a). **c)** Arrangement of selection regions considered in this work: **i**: the ‘isolated’ scenario, where a selection region (dark gray) is embedded in an infinitely large backdrop of refuge (white), **ii**: ‘infinite periodic’ scenario of infinitely many selection and refuge regions, **iii**: ‘multiple periodic’ scenario of multiple selection regions separated by refuge regions embedded in an infinite backdrop of refuge.

### Model

Motivated by agricultural insecticide application, we introduce a minimal one-dimensional model that qualitatively demonstrates the evolutionary consequences of a spatio-temporally heterogeneous measure to control a motile population. To facilitate applications beyond evolution of resistance to pesticides, we will mostly use language agnostic to the evolving system. In the model, formally described in Section S1: Mathematical Model Description, the population exhibits standing genetic variation as well as undirected motility and breeds once a year. We consider two genotypes that map to phenotypes with respect to action of the control agent. Population *S* experiences the effects of a control agent; its spatial distribution is described by density *u*_*S*_(*x, t*) at location *x* and time *t*. Population *R* is resistant to the control agent and analogously described by *u*_*R*_(*x, t*). As a simplifying assumption, to be discussed later, we consider both densities to be uniform at the start of the generation with *u*_*R*_(*x, t* = 0) = *ϕ* and *u*_*S*_(*x, t* = 0) = 1 *− ϕ* assuming a dimensionless carrying capacity of 1.

We model motility and consequently dispersal by diffusion affecting both populations independently (notation *u*_*S,R*_)):

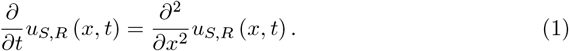

*t* is time in units of generation time *Y*, i.e., the fraction of generation time passed; we here only consider 0 *≤ t ≤* 1. Similarly, *x* is location along the axis measured in units of typical dispersal distance within one generation, 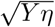 with *η* the unscaled diffusion coefficient. Note that we here ignore that insect dispersal depends on life stage and choose diffusion as simplest representation for continuous dispersal (as previously employed in Ref. [Com77]).

Space is partitioned into two types of regions: We refer to regions where the control agent is applied as ‘selection regions’ (gray in Fig. 1a) and regions in which they are never applied as ‘refuge region’ (white in Fig. 1a), corresponding to another species of crop, an untreated cropping region, or the natural environment. Note that presence of control measure is the only difference between selection and refuge regions, e.g., there is no difference of dispersal within those regions and no barriers of dispersal between them.

The control agent is applied to the selection regions from the start for part of a generation time, i.e., for *t* with 0 *< t* ≤ *p* with *p <* 1. During this time, any susceptible individuals within the selection regions are removed, whilst resistant individuals are unaffected. This corresponds to *u*_*S*_(*x, t*) = 0 inside selection regions and an absorbing boundary condition at the edges of the selection region, affecting the density of the susceptible population around the selection region up to a distance of 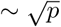; see Fig. 1a for *t* = 0 and for *t* = *p*. For *p < t <* 1, the control measure is not applied, and diffusion leads to smoothing of the density profile as seen in Fig. 1a for *t* = 1 just before breeding takes place. This results in an influx of population S into the previously controlled region at times *p < t <*1, up to a distance of 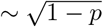. Note the emergence of two length scales, 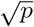 and 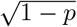. If *p >* 0.01, i.e., the control measure is applied for a reasonable part of the generation time, both length scales are in the range of 0.1 to 1.

Breeding is modeled as taking place at *t* = 1 via a normalization procedure: the total density of both populations is increased to the carrying capacity (set to 1) at each point in space, with the relative proportions of each population maintained. Note that no de novo mutations for resistance are assumed to occur during breeding. Denoting *U*_*S,R*_(*x*) as population density of the susceptible and resistant population after breeding at *t* = 1, respectively, this can be formalized as

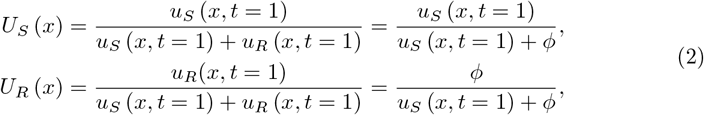

where we understand *u*_*S,R*_(*x, t* = 1) as representing population time before breeding and have taken into account that *u*_*R*_ is unaffected by the control measure and due to the uniform initial condition unchanged throughout the season, i.e., *u*_*R*_(*x, t*) = *ϕ*. The densities after breeding are visualized in Fig. 1a at *t* = 1 after the breeding phase (note the log scale for population density).

We quantify the effect of the control agent on population composition over a generation by the change Δ*R* in the number of resistant individuals through breeding:

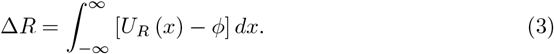

This change Δ*R* depends on the size and distribution of selection regions and is the core observable of this study. A rise in the number of resistant individuals is due to selection imposed by the control measure. To account for the fact that the control measure has benefits, we normalize Δ*R* by the size of the region in which the control measure is applied; in agriculture this would relate to the rise in resistant insects per size of farmland protected.

Of the vast number of possible different distributions of selection regions, we identified three prototypical examples to focus on:

#### i Isolated

A selection region of size *w* is embedded in an infinitely large backdrop of refuge (Fig. 1a and Fig. 1ci).

#### ii Infinite periodic

The landscape consists of infinitely many selection regions of width *w* and neighboring refuge regions of width *d* (Fig. 1cii).

#### iii Multiple periodic

A finite number *N* of selection regions of size *w* are separated by refuge regions of size *d* embedded in an infinite backdrop of refuge (Fig. 1ciii).

## Results

### Isolated Selection Region

We first consider the scenario of an isolated selection region. To find the normalized change in the number of resistant individuals for an isolated selection region, Δ*R*_*I*_ (*w*)*/w*, for parameters *w* (selection region size), *p* (fraction of generation for which control agent is applied) and *ϕ* (frequency of resistant phenotype at start), we need to solve the diffusion equation (Eq. 1) within refuge regions with Dirichlet boundary conditions: *u*_*S*_(*x, t*) = 0 for *x* = 0 and *x* = *w* as well *u*_*S*_(*±*∞, *t*) = 1 *− ϕ*. As detailed in Section S2: Isolated, this can be achieved by Laplace transform and contour integration. The solution for *t* = *p* serves as initial condition when solving the diffusion equation in the whole domain for *t > p* by numerical convolution of *u*_*S*_(*x, p*) with the diffusion kernel. Numerical evaluation of Eq. 2, representing breeding, and integration over space (Eq. 3) then provides the change in the number of resistant individuals per width of selection region, Δ*R*_*I*_ (*w*)*/w*. Figs. 1b and 2a display Δ*R*_*I*_ (*w*)*/w* for *p* = 0.1 and *ϕ* = 10^−5^, which displays a pronounced minimum at *w*^*^ ∼ 1.

The existence of a global minimum can be intuited by considering limiting values of *w*. For *w* = 0, the removal of susceptible individuals at a point during the selection phase leads to a decrease in *u*_*S*_(*x, t*). After breeding, application of the control measure at a single point has therefore caused a finite increase in the number of resistant individuals, lim_*w*→0_ Δ*R*_*I*_ (*w*) → Δ*R*_*I*,0_ *>* 0, and thus lim_*w*→0_ Δ*R*_*I*_ (*w*)*/w* → ∞. For 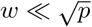 and 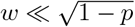, we expect Δ*R*_*I*_ (*w*) ≈ *R*_*I*,0_ and thus Δ*R*_*I*_ (*w*)*/w* ∼ *w*^−1^, indicated as a dashed green line in Fig. 2a and matching the numerical solution well.

**Fig 2.**
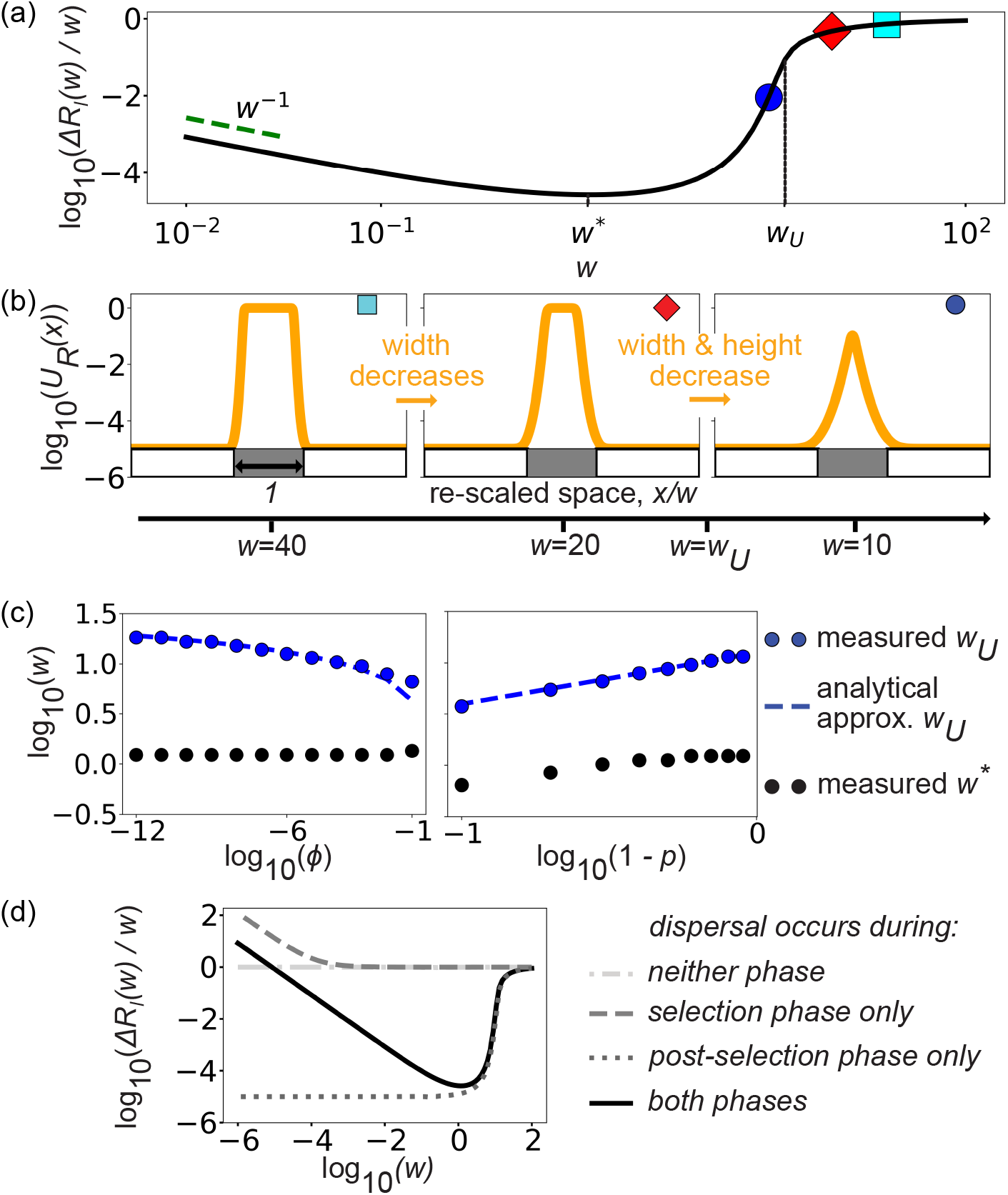
Characterization of the increase in the resistant population normalized by size of the isolated selection region. *p* = 0.1, *ϕ* = 10^−5^ unless otherwise specified. **a)** Δ*R*_*I*_ (*w*)*/w* exhibits a minimum, bounded by a regime where ∼ *w*^−1^ scaling for small *w* (green line) and a region with rapid change in slope at *w*_*U*_ (indicated). Symbols refer to the panels in **b)**, which displays *U*_*R*_(*x*), the density of the resistant population after breeding for different *w*. As the width *w* of the selection region decreases, the density profiles of susceptible individuals at *p* = 1 are able to move closer together until they overlap at around *w*_*U*_, at which point the peak of *U*_*R*_(*x*) shrinks. Note that for visualization purposes, *x* got rescaled by *w*. **c)** Measured *w*_*U*_ (blue dotted), analytically derived *w*_*U*_ (blue dashed, Eqn.4), and measured global minimum *w*^*^ (black dotted) for changing *ϕ* (left, *p* = 0.1) and changing *p* (right, *ϕ* = 10^−5^). **d)** Δ*R*_*I*_ (*w*)*/w* for different scenarios of dispersal: no dispersal [gray dash-dotted line], only during selection phase [gray dashed line], only after selection phase [gray dots], or during both phases [black line]).

Conversely, for sufficiently large *w*, susceptible individuals only diffuse a distance up to 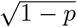 into the selection region on either side after selection. Thus, for 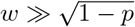, the whole region apart from its boundaries is void of susceptible individuals before breeding. Thus, the density profile of the resistant phenotype is plateau-shaped at a large scale (*U*_*R*_(*x*) ≈ 1 inside the selection region, Fig. 2b, cyan square) and thus Δ*R*_*I*_ */w*∼ (1 *− ϕ*). The effect of boundaries declines with increasing *w*, and thus is approached from below. This completes the quantitative justification for the global minimum: Δ*R*_*I*_ (*w*)*/w* is a strictly decreasing function for *w* ≪ 1 and a strictly increasing function for *w* ≫ 1.

Combining the considerations for small and large *w* and using that 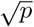 and 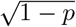 are within an order of magnitude from 1, we expect a global minimum *w*^*^ around 1. The exact location *w*^*^ depends on both the initial proportion of resistant individuals, *ϕ*, and the proportion of the generation over which the control measure is active, *p*, depicted in Fig. 2c. While we have no analytical prediction for the precise location of the global minimum, *w*^*^, we can derive an upper bound, *w*_*U*_. Fig. 2a,b illustrate how Δ*R*_*I*_ (*w*)*/w* decreases as the plateau of *U*_*R*_(*x*) shrinks and disappears. We expect a significant change in Δ*R*_*I*_ (*w*)*/w* for decreasing *w* when susceptible and resistant population densities are equal in the center of the selection region (*x* = *w/*2). As *ϕ* ≪ 1, we expect this to occur at relatively large *w*, at which point the constant-size boundary effects during the selection phase have a smaller proportional effect and are therefore ignored. In this approximation, *u*_*S*_(*x, p*) ≪ 1 inside the selection region and 1 − *ϕ* outside at the end of the selection phase. The distribution of susceptible individuals before breeding, *u*_*S*_(*x* = *w/*2, *t* = 1) can be found by convolution with the diffusion kernel (see S3: *w*_*U*_ Derivation):

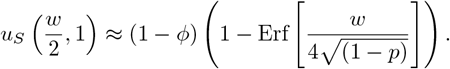

Demanding *u*_*S*_(*x* = *w/*2, 1) = *u*_*R*_(*x* = *w/*2, 1) = *ϕ*, we obtain

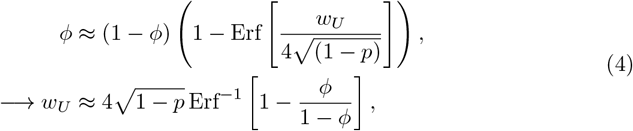

where the scaling with 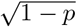 can be intuited by diffusion from the boundary over time 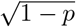. Fig. 2c also indicates the analytically derived *w*_*U*_ obtained above, which agrees well with the location of the fastest change in slope (the measured *w*_*U*_).

### Factors driving growth of resistant population

Above, we reasoned the existence of a minimum in Δ*R*_*I*_ (*w*)*/w* based on limiting behavior for small and large *w*. We will now describe the role of dispersal in the emergence of the minimum. We start by considering the three key factors of this model which modulate the post-breeding distribution of the resistant population:

i. Firstly, susceptible individuals are lost due to the blanket implementation of selection pressure at *t* = 0. This effect removes (1 *− ϕ*)*w* from the susceptible population immediately. In the absence of dispersal, this results in Δ*R*_*I*_ (*w*)*/w* = 1 *− ϕ* for all *w* (light gray dash-dotted line in Fig. 2d).
ii. Secondly, dispersal during the selection phase acts to reduce the susceptible population in a ‘boundary region’ of width 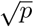 in the vicinity of the selection region (independent of the width of the selection region). By considering the case where dispersal only occurs during the selection phase (dark gray dashed line in Fig. 2d), we note that the effect of this dispersal only significantly affects Δ*R*_*I*_ (*w*)*/w* for 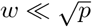, i.e. when the amount of the susceptible population lost in the selection phase is comparable to the amount lost due to the blanket application of selection pressure.
iii. Lastly, dispersal during the post-selection phase enables the loss of the susceptible population to be more evenly distributed over space: The first impact is a recolonization of the selection region (which would act to decrease Δ*R*_*I*_ (*w*)*/w*), with a secondary impact being a decrease in the overall susceptible population beyond the boundary layer up to a distance ≈ 1 compared to the initial susceptible population distribution (which would act to increase Δ*R*_*I*_ (*w*)*/w*). Recolonization is the primary factor by which the post-breeding size of the resistant population is modulated to be smaller than in the absence of any dispersal: This is evidenced by Δ*R*_*I*_ (*w*)*/w* being smaller in the case of migration only occurring during the post-selection phase (dark gray dotted line in Fig. 2d) compared to the case of no-migration (light gray dash-dotted line in the same figure) for all *w*.

Note that the role of factor (iii) to decrease Δ*R*_*I*_ (*w*)*/w* is impacted by both factors (i) (the relative proportion of the selection region that can be recolonized shrinks as *w* increases) and (ii) (the boundary layer acts to decrease the amount of susceptible population available to recolonize the selection region, reducing the effective penetration depth). It is in these competing effects that we see that the dynamic of a global minimum in Δ*R*_*I*_ (*w*)*/w* can only transpire when dispersal occurs both during and after the selection phase (black line in Fig. 2d). Therefore, we interpret the global minimum to be due to a trade-off between the loss of susceptible individuals in the vicinity of the selection region during the selection phase, and the subsequent recolonization of the selection region during the post-selection phase.

### Periodic Selection Regions

We next consider an infinite periodic sequence of staggered selection regions of width *w* and refuge region of width *d* (Fig. 1cii). The natural observable for measuring growth of the resistant population is growth in a single sub-unit, denoted by Δ*R*_*P*_ and obtained from Eq. 3 when limiting integration to a sub-unit. Δ*R*_*P*_ now depends on both the width of the selection region, *w*, and the separation between selection regions, *d*. For *d* ≫ 1, the individual selection regions are effectively independent and emergence of the resistant population matches that of the isolated region discussed above.

Δ*R*_*P*_ can be obtained using a combination of analytical and numerical methods. Solving the diffusion equation (Eq. 1) with boundary condition of *u*_*S*_ vanishing inside the selection regions results in an analytical expression for the density of sensitive individuals for 0 *< x < d* (using the method of separation of variables, see Ref. [RHB06]):

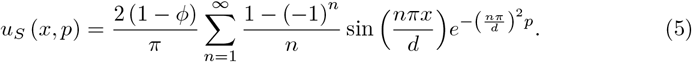

From here we can solve the diffusion equation for one subunit subject to periodic boundary conditions using Eqn. 5 as the initial condition to analytically evaluate the pre-breeding distribution of susceptible individuals (see S4: Periodic). Following normalization to simulate the breeding process, integration provides us with Δ*R*_*P*_.

Motivated by the question on how to minimize emergence of the resistant population given a fixed ratio of selection region and refuge, we consider a new set of variables: *k* = *w* + *d*, the size of a sub-region, and *α* = *d/k*, the fraction of the sub-region which is refuge. Fig. 3a displays the Δ*R*_*P*_ (*k*)*/w* for continuous *k* and a set of different *α*. Keeping the ratio of refuge and selection region fixed (fixed *α*) we observe a minimum bounded by two plateaus for small and large *k*. In the following, we will rationalize the plateaus and determine bounds for the location of the minimum.

**Fig 3.**
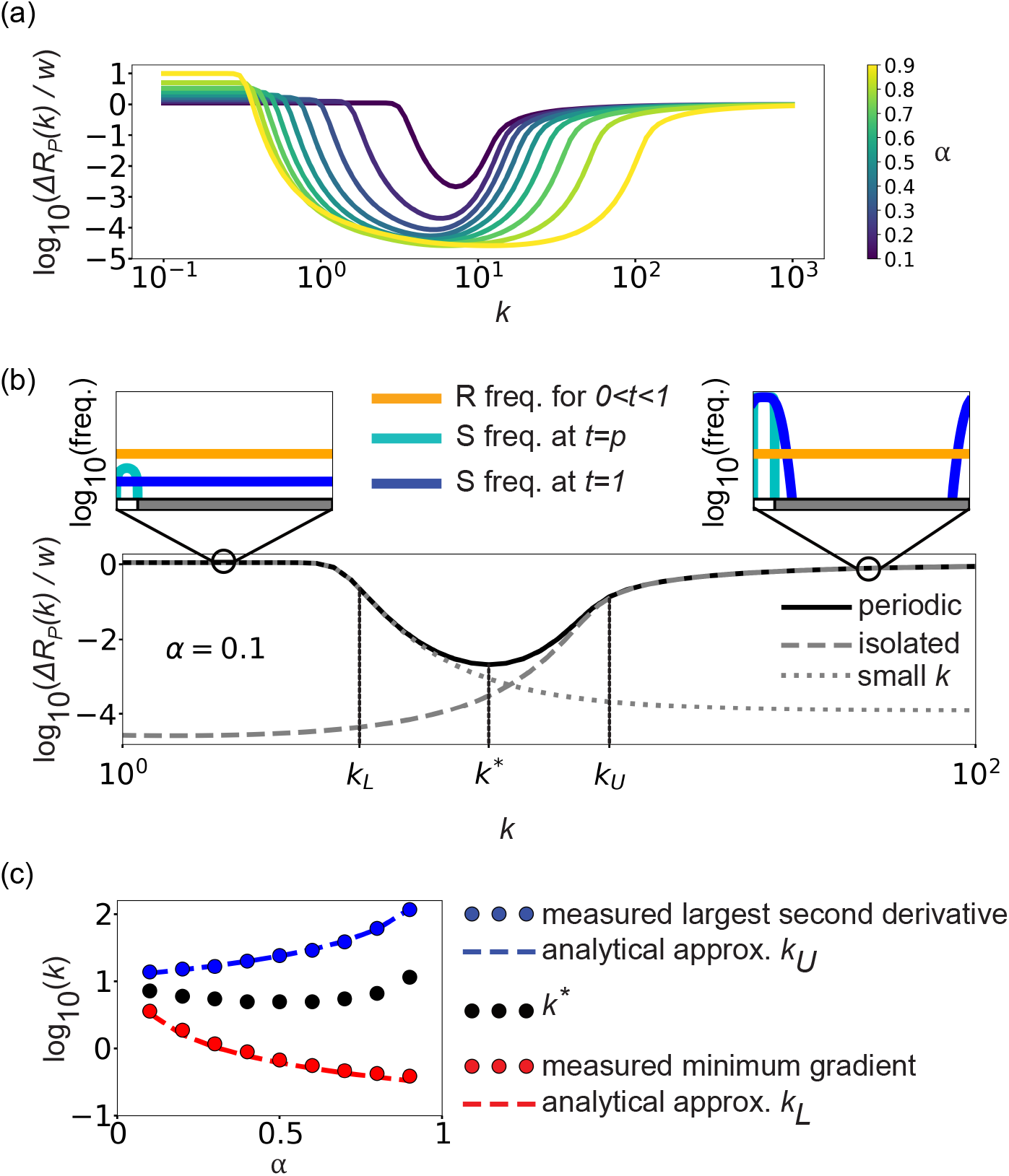
Characterization of the increase in the resistant population normalized by size of the selection region for a periodic arrangement of selection regions. **a)** Change in the number of resistant individuals, Δ*R*_*P*_ (*k*)*/w*, over a periodic sub-unit per width of the selection region, *w*, for a variety of values of the proportion of a periodic sub-unit of refuge region, *α*, for *p* = 0.1 and *ϕ* = 10^−5^, with a pronounced minimum for all *α*. **b)** Δ*R*_*P*_ (*k*)*/w* for *α* = 0.1 (black line) with location of steepest slope and largest change in slope (obtained numerically) indicated at *k*_*L*_ and *k*_*U*_, respectively. For large *k*, Δ*R*_*P*_ (*k*)*/w* is well-described by Δ*R*_*I*_ (*w*)*/w* (dashed line) and for small *k* by Eqn. 7 (dotted line). **c)** Location of the minimum *k*^*^ together with numerically (red and blue dots) and approximate analytically determined bounds (red and blue dashed lines, Eqs. 8 and Eq. 6, respectively) as a function of *α*.

Let us first focus on the case of sufficiently large *k*. As stated above, for large refuge regions, *d* = *k · α* ≫ 1, growth of the resistant population is well described by considering isolated regions: Δ*R*_*P*_ (*k*)*/w* (black line) and Δ*R*_*I*_ (*w*)*/w* (dashed line) in Fig. 3b overlap for *k > k*_*U*_. Consequently, we follow the arguments for the isolated region to find an upper bound for the location of the minimum at *k*^*^ and obtain

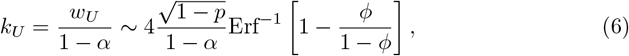

which matches well the location of largest change in slope (Fig. 3c, blue).

For small *k, k* ≪ 1, the individual selection regions influence each other as diffusion is fast enough to for neighboring selection regions to interact. In this regime, the first term in Eq. 5 dominates due to the exponential decay:

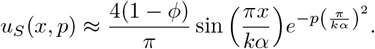

Let us assume that the periodic sub-unit is sufficiently small such that the susceptible individual population *u*_*S*_(*x, t*) flattens out by *t* = 1, i.e.,

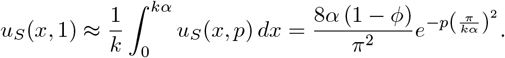

Consequently, for small *k*, after breeding and integration over space,

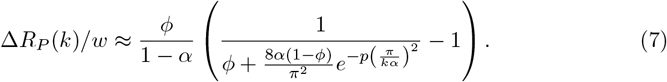

This is a monotonically decreasing function of *k*, indicated as a dotted line in Fig. 3b, which, for chosen *α* = 0.1, *p* = 0.1, and *ϕ* = 10^−5^, follows the shape of Δ*R*_*P*_ (*k*)*/w* up to *k* ∼ 10^0.5^. We define *k*_*L*_, the lower bound for the position of the minimum, as the *k* for which Δ*R*_*P*_ (*k*)*/w* has the steepest negative slope. Using the small system considerations, this point can be computed analytically from Eq. 7 as

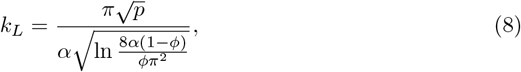

which agrees very well with the numerically determined *k*_*L*_ of the full solution (Fig. 3c).

We discussed above that emergence of resistance is suppressed by diffusion of the susceptible population into the selection region after selection. This is not possible for large selection regions (in both the isolated and periodic cases) because the population is unable to penetrate deeply into the selection region. For periodic landscapes, this is also not possible if the refuge is too small, as in this case the susceptible population is diminished too strongly during the selection step; note the exponential decay with decreasing refuge region width *d* in Eqn. 5. For fixed *α*, the minimum thus emerges between small and large *k*.

### Multiple Selection Regions

Minimizing the emergence of resistance is subject to constraints both in aim and practicality. For example, to acknowledge the need to control pests in the first place, we normalized growth of the resistant population by the size of the region affected by the control measure. Similarly, one may want to control a given total amount of landscape, but may have options on how to place individual sections or fields. To address these possible constraints, we consider a large selection region of width *c* sub-divided into *N* smaller regions of width *w* and consider Δ*R/c*, the change in the resistant population normalized by the overall size of the selection region. We consider two cases: First, sub-regions may be separated arbitrarily far apart; this implicitly assumes that the selection regions have no limits on where they can be placed, so we term this the ‘unbound space’ scenario (Fig. 4a). Second, we introduce an additional constraint by specifying the finite total amount of space where it is possible for the selection region to be placed, of size *h*. This case is referred to as the ‘bound space’ scenario (Fig. 4b).

**Fig 4.**
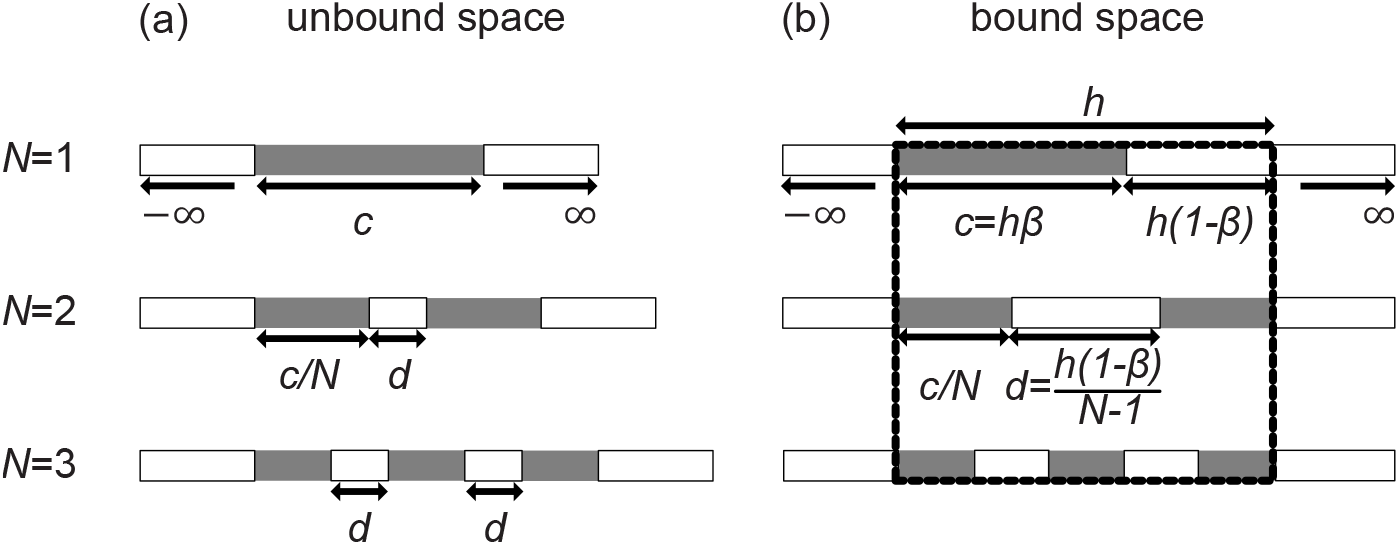
Examples of each of the types of multiple selection region distributions. The ensemble of selection regions (gray) is surrounded by an infinitely large refuge region (white). **a)** The ‘unbound space’ scenario, where a total selection region width of *c* is split into *N* regions with constant separation, *d*, between them. **b)** The ‘bound space’ scenario, where there is a total region of width *h* (bound by dashed lines) within which the selection region is constrained. A proportion *β* of this region is selection region, so the total selection region width is *c* = *h* ·*β* and the enclosed refuge region is *h* · (1 − *β*). For *N* selection regions, the width of the separation between selection regions is therefore *d* = *h ·* (1 *− β*)*/*(*N −* 1) for *N >* 1.

To determine the growth of the resistant population, we are using a combination of analytical and numerical approaches as above, described formally in S5: Multiple. In short, *u*_*S*_(*x, p*) is obtained using Eq. 14 (where the refuge extends to infinity to the left and right of the system) and Eq. 5 (when the refuge is wedged between two selection regions such as in the periodic case). Convolution with the diffusion kernel yields *u*_*S*_(*x*, 1) and integration (to simulate breeding) results in Δ*R*, from which we obtain the observable of interest, Δ*R/c*.

For particular regimes, we expect the rise in the resistant population to be well-described by approximations to the scenarios considered previously:

- The ‘individual isolated approximation’ considers the landscape as a finite number of completely isolated regions of width *w* = *c/N* and thus Δ*R/c* ≈ Δ*R*_*I*_ (*w*)*/w*. We expect the approximation to be valid for *d* ≫ 1, i.e., when the selection regions are far enough away such that dispersal by diffusion is unable to cover the distance between selection regions, so selection regions can be considered to be independent.
- In the ‘periodic approximation’, we disregard the refuge regions to the left and right and instead consider *N* sub-units with periodic boundary conditions. Here, the overall change in resistance is the product of *N* and the change in resistance per periodic sub-unit (of width *k*) selection region width, Δ*R/c* ≈ *R*_*P*_ (*k*)*/w*, where again *w* = *c/N*. We expect this approximation to be valid in two regimes: First, when the overall width of the sub-units is large enough such that the effects of the infinitely large refuge regions at either end penetrate only a relatively small distance into the bulk of the region. Second, when the size of the interspersed refuge regions is sufficiently large such that there is little to distinguish between them and the infinite regions of refuge: In this case, this approximation matches the ‘individual isolated approximation’ described above.
- Finally, the ‘combined isolated approximation’ assumes that the entire space between the refuges on the left and right approximates the behavior of a single selection region. The overall change in resistance is calculated identically to a single isolated region of width *w*: Δ*R/c* ≈ Δ*R*_*I*_ (*w*)*/w*, where *w* = *c* + (*N* − 1)*d* (for the unbound case) or *w* = *h* (for the bound case when *N >* 1; for *N* = 1: *w* = *hβ*).

We expect this approximation to be valid if the separation between selection regions is small 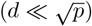 such that by the end of the selection phase there is a negligible number of susceptible individuals left in the interspersed refuge regions.

### Unbound Space

Fig. 5 shows the effect of subdivision for the case of ‘unbound space’, i.e., the number of selection regions *N*, on growth of the resistant population, specifically Δ*R/c*, for given overall size of the selection region, *c*, and size of refuge regions, *d*. In each case, we see overall agreement with the hypotheses listed above:

**Fig 5.**
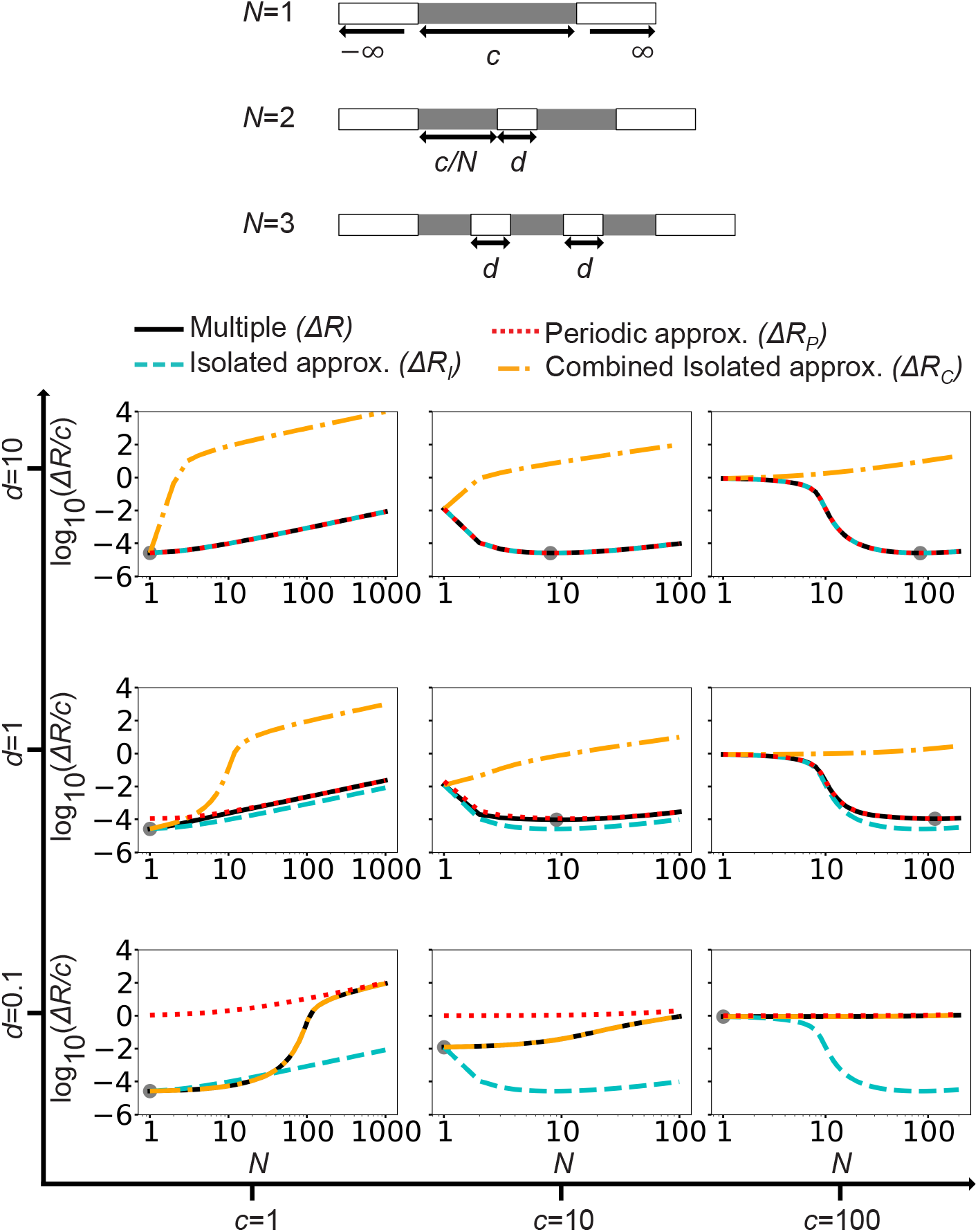
A comparison of the change in resistant individual number for the unbound case of multiple selection regions (black solid line), compared to approximations constructed from *N* isolated regions (cyan, dashed line), *N* periodic sub-units (red, dotted line) and one large isolated region (orange, dash-dotted line), shown for a variety of total selection region widths, *c*, and separation between selection regions, *d*, highlighting how (*c, d*) values determine the applicability of the approximations. In each case, *p* = 0.1 and *ϕ* = 10^−5^. The global minimum of unbound case of multiple selection regions is indicated by a gray dot.

- The ‘individual isolated approximation’ (cyan dashed line) is an accurate description of the multiple selection region case for large refuge region (*d* = 10, top row), as in this regime the selection regions are far enough to be independent over the course of a single generation. Note that for *d* = 1, 0.1, the individual isolated approximation is accurate only for *N* = 1 (when the multiple case is identical to an individual isolated case).
- The ‘periodic approximation’ (red dotted line) is an accurate description for large refuge regions (*d* = 10, top row), corroborating our second hypothesis that this approximation is accurate in the regime where the infinitely large refuge regions approximate the behavior of one of the large interspersed regions. For intermediate refuge regions (*d* = 1) with small overall selection region width (*c* = 1), the periodic approximation only becomes accurate when *N* ⪆ 10, while being relatively accurate for all *N* for *d* = 1, *c* = 10, 100. For small refuge region size (*d* = 0.1), for *c* = 1 and *c* = 10 there is a marked discrepancy between the multiple case results and the periodic approximation, until *N* ≈ 500 and *N* ≈ 100, respectively. For (*c, d*) = (10, 1), (100, 0.1) and (100, 1) the periodic approximation is valid for all *N*. All of these scenarios correspond to large overall system sizes where the boundary effects would be able to penetrate a relatively short distance into, thus the accuracy of the periodic approximation agrees with our second hypothesis. Specifically, these results confirm the accuracy of this approximation when *c* + (*N −* 1)*d* ≫ 1.
- The ‘combined isolated approximation’ (orange dash-dotted line) is, as expected, a very accurate result in the regime where *d* is small (*d* = 0.1), as here there are few surviving susceptible individuals remaining by the end of the selection phase, such that the overall distribution of the susceptible population is similar to a large selection region of size *c* + (*N −* 1)*d*. This observation is in agreement with our third hypothesis.

The location of the global minimum of the full system agrees with that of the most suitable approximation. For *c* = 1 or *d* = 0.1 we note that the global minimum corresponds to *N* = 1. For *c* = 1 this can be understood by recalling that both the isolated and periodic cases predict an optimum to occur for *w* ≈ 1: Increasing *N* in this regime has the effect of decreasing the width of the selection regions to smaller values, therefore *N* = 1 would correspond to the smallest attainable value. In the case of *d* = 0.1, this can be understood by realizing that the ‘combined isolated approximation’ is the most accurate: As this approximation assumes that the effective selection region size is *w* = *c* + (*N −* 1)*d*, similar to the isolated case we see that the expected optimum of *w* ≈ 1 is closest when *N* = 1 in all cases. For the remaining examples, note that the minimum occurs at *N* ≈ *c*, i.e., *w* ≈ 1 as remarked earlier.

### Bound Space

While the ‘unbound space’ scenario sheds lights on the trade-offs of subdividing a selection region across an arbitrarily large region, it fails to consider the constraint that all selection may need to occur in a finite space due to geographical or ownership constraints. The ‘bound space’ scenario can take these constraints into account. When subdividing the selection region further, both individual selection and refuge regions become smaller.

Fig. 6 shows the effect of subdivision for the ‘bound case’, i.e., the number of selection regions *N*, on growth of the resistant population, specifically Δ*R/c* for a given fraction of selection region, *β* and total size *h*. We observe a clear minimum as a function of the number of individual regions, *N* (black line). For all data presented, the optimal subdivision corresponds to a selection region size of *w* ∼ 1, as has been previously reasoned in the isolated case.

**Fig 6.**
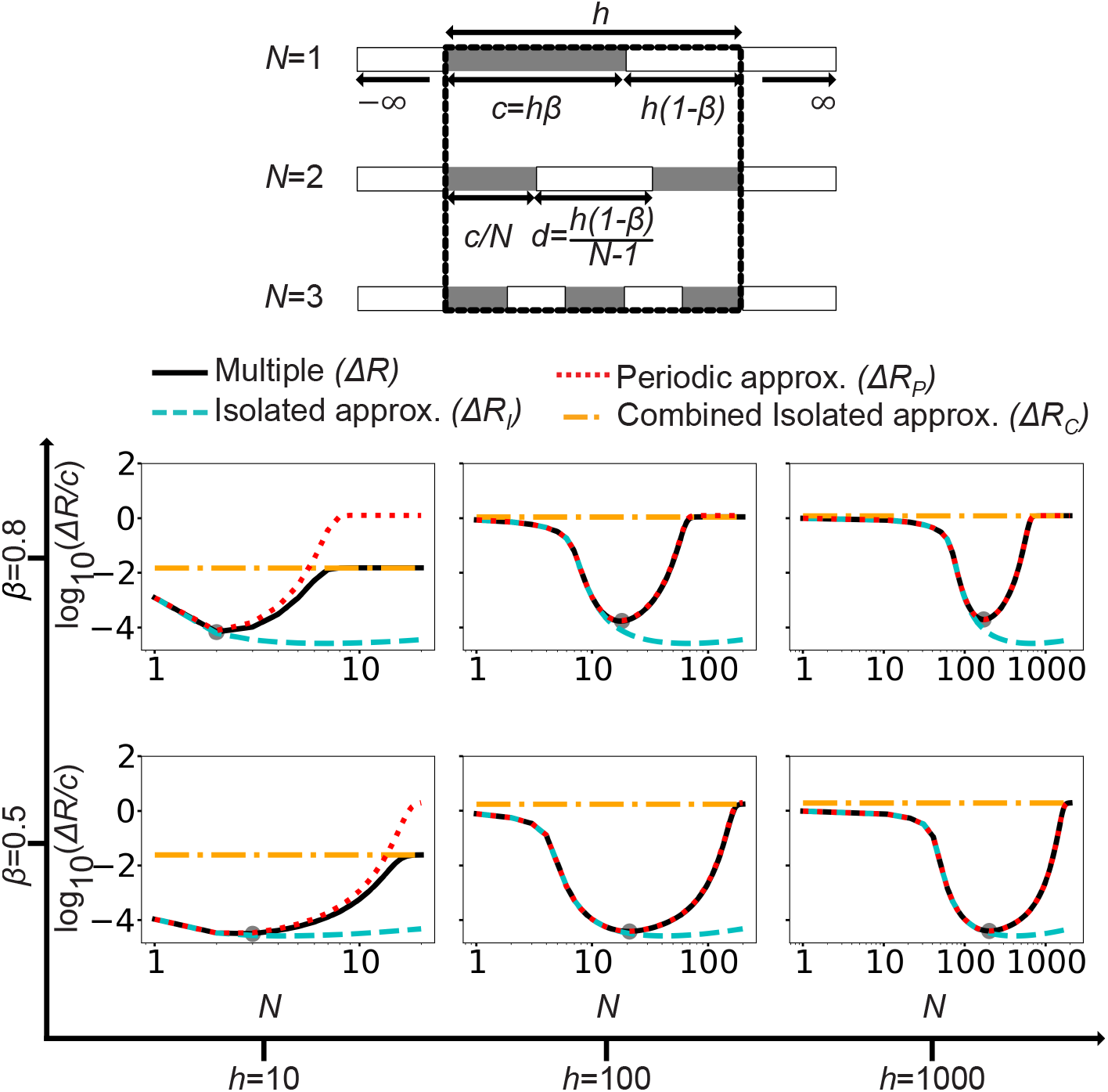
A comparison of the change in the number of resistant individuals for the bound case of multiple selection regions (black solid line) compared to approximations constructed from *N* isolated regions (cyan, dashed), *N* periodic sub-units (red, dotted) and one large isolated region (orange, dash-dot), shown for a variety of total bound region widths, *h*, and the proportion of the region that is selection region, *β*, highlighting how (*h, β*) values determine the applicability of the approximations. In each case, *p* = 0.1 and *ϕ* = 10^−5^. The global minimum of the bound case of multiple selection regions is indicated by a gray dot.

Aspects of the overall shape are again well described by the different approximations discussed above.

- The ‘individual isolated approximation’ (cyan dashed line) is accurate for all *N* smaller than which corresponds to the global minimum in the isolated case. This is because the global minimum occurs when *w* ∼ 1 for the isolated case, and this corresponds to when *d* is of the same order of magnitude in the values of *β* presented in Fig. 6. Therefore, increasing *N* beyond the optimum acts to shrink the interspersed selection regions such that the selection regions are no longer effectively independent.
- The ‘periodic approximation’ (red dotted line) is accurate for all *N* when the overall system is large (*h* ≥ 100), in that case edge effects penetrate a relatively small region into the ensemble of selection regions. In contrast, for small *h* (*h* = 10), we note that this approximation becomes more inaccurate with *N* increasing: for small *N* the infinite refuge regions approximate the behavior of the interspersed selection regions, but for larger *N* this stops being the case and the resultant edge effects perturb the system further away from the periodic result.
- The ‘combined isolated approximation’ (orange dash-dot) acts as the limiting behavior for large *N* relative to the system (such that *N* ∼ *h*). It is in this regime that the size of the interspersed refuge regions (*d* = *h*(1 − *β*)*/*(*N* −1)) is sufficiently small such that few susceptible individuals remain within them by the end of the selection phase, resulting in an effective isolated selection region of width *h*.

In both the bound and unbound scenarios, we see that the isolated and periodic cases can be taken as accurate approximations in predictable regimes, as laid out in our hypotheses and later validated by direct comparison with simulations. More generally, the bounds ascertained in the isolated and periodic sections can be employed as quick heuristics to assess the efficacy of a given selection region distribution in minimizing the rate of resistance emergence. Both Fig. 5 and Fig. 6 act as phase diagrams highlighting the relative accuracy of each approximation for each spatial constraint (for the unbound and bound scenarios, respectively).

## Discussion

Using a minimal model of resistance evolution which captures continuous spatial and temporal features of heterogeneous landscapes, we have demonstrated that the distribution of selection pressure can be tuned to minimize the initial rate of rise in resistance in the population, in cases where there are no spatial limits in distribution (Fig. 5) and when there are such limits (Fig. 6). We can qualitatively understand the emergence of this optimum by considering the dynamics (loss and dispersal) of the susceptible population throughout a generation. Further simplified scenarios (‘isolated case’ and ‘periodic case’) allow us to predict the location of the optimum for particular regimes of a given distribution.

We highlighted that the emergent dynamics associated with an optimal distribution of selection pressure arise only when continuous population dispersal takes place in an environment in which the selection pressure is both spatially and temporally heterogeneous. Our model is thus truly minimal in the sense of capturing the essential features leading to a minimum in rise of resistance. This is in contrast to previous models which have further simplified dispersal and spatio-temporal heterogeneity: For example, previous work also employed a continuous diffusive dispersal behavior of pests [Com77], yet neglected the dynamics of dispersal of susceptible individuals into the selection region, which we have demonstrated to dramatically impact the rate of resistance increase (Fig. 2d). This model also explores a gap in the literature: Having no fitness cost leads to unbounded rise in resistance over time, which in the literature motivates research into the trade-off between a fitness cost and separation between selection regions [Mui75; LR98]. While the rise of resistance in the absence of a fitness cost may be inevitable, the rate at which it rises can be tuned and minimized, which we have demonstrated.

The model considered here is minimal by design and needs to be extended to capture the dynamics of a pest and its control in nature. This includes dispersal, here modeled as diffusive, which in nature may include long-distance jumps and targeted movement (such as in Ref. [Str+14]). We furthermore assumed full effectiveness of the control measure on susceptible individuals and full resistance to the control measure by resistant individuals, whereas in a real-world system, the efficacy of a pesticide may be impacted by the dose the individual has acquired. While we expect that these additional features will impact the optimal distribution of control measure, the underlying process of selection followed by recolonization ought to preserve the existence of such an optimum.

We also studied a one-dimensional system, while two-dimensional domains are relevant for applications. However, we expect the findings to be easily extended to two-dimensional systems. In the case of an isolated rectangular field, we anticipate that the narrowest dimension will take the place of *w*, as this will be the dimension over which there will be a deeper recolonization relative to the width of the dimension. In the case of a sequence of rectangular selection regions, this is in many ways analogous to a one-dimensional system with the multiple case. When field shapes and distributions are more complex, we are confident that the qualitative and quantitative justifications for the existence of an optimized distribution will still hold.

Last, but not least, we focused on the initial rise in resistance assuming uniform initial conditions, in both carrying capacity and population distribution. This situation only realistically occurs in scenarios where a new cropping region is set up (thus previous years of farming have not been able to significantly affect the spatial distribution of pests) and resistance exists on a background level only due to stochastic effects (for example, due to rare de novo mutations, or perhaps due to migration from regions where insecticide has been applied). Even if these conditions are not fulfilled, the qualitative and quantitative reasoning for the emergence of an optimized selection region distribution remain the same, thus we hypothesize that even with non-uniform initial conditions there will still be a global optimum in relative change in resistant individual frequency, albeit perturbed from the results shown here.

Throughout this work have focused on how to minimize the initial rate of resistance emergence on the local scale with a single variety of control measure: This suggests three natural avenues for further research. The first is to consider the optimal distribution of control measure over many generations: This may differ from the optimal distribution for one year as a result of the change in sub-population distribution at the start of each generation. Furthermore, would necessitate considering the occurrence of de novo mutations if a large number of generations are considered. The second is to consider how optimization on the local scale can tie into existing research considering control measure distribution on the geographical scale. For example, in the presence of a pest population undergoing range expansion, how the distribution of the selection pressure and the fitness cost to resistance interact to shape the genetic composition of the population frontier has been determined at a coarse-grained scale [TRM23]. Such a multi-scale approach to selection pressure application could create a robust distribution of control measure which best minimizes the frequency of resistant individuals across a large area. Thirdly, this modeling framework can be extended to determine how a variety of different control measures (effective on different genotypes) can be best distributed to minimize the overall rate of resistance evolution to each control measure.

In summary, the rate of resistance evolution can be minimized by carefully tuning the selection pressure distribution in time and space. Reducing this rate will lead to a longer increased lifetime in control measure application, allowing more time for the production and testing of novel control measures. We hope that this work will therefore inform policy makers of top-down distribution of control measures. For example, how individual farmers or groups of farmers may stagger pesticide application in space to minimize resistance evolution.

## Acknowledgments

Thomas Tunstall acknowledges support by EPSRC DTP and Syngenta Crop Protection. Wolfram Möbius acknowledges support by BBSRC via BBSRC-NSF/BIO grant BB/V011464/1.

## Supporting information

### S1: Mathematical Model Description

#### Initial Conditions: *t* = 0

In the instant before control measure application, the population is uniformly distributed over all space. We choose the population density to be normalized to 1, which we will take to be the carrying capacity of the total population. The total proportion of resistant individuals is *ϕ* : *ϕ* ∈ [0, 1], therefore the total proportion of susceptible individuals is 1 − *ϕ*, everywhere.

At *t* = 0, the control agent is applied within the selection regions: any susceptible individuals within this region are removed, whilst the resistant ones are completely unaffected. Formally, the initial conditions are:

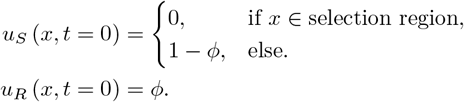

These distributions are illustrated in Fig. 1a (*t* = 0).

#### 1) Selection Phase: 0 *< t ≤ p*

The control measure is active for a proportion *p* of the generation, *p* ∈ (0, 1): during this time, any susceptible individuals that diffuse into the selection regions are immediately removed. This is akin to Dirichlet boundary conditions being applied to the edges of the selection region, ∂_Selection_, for the susceptible population density, *u*_*S*_(*x, t*), only. As the resistant individuals are not affected by control agent, the density profile of the resistant population, *u*_*R*_(*x, t*), remains constant. Formally:

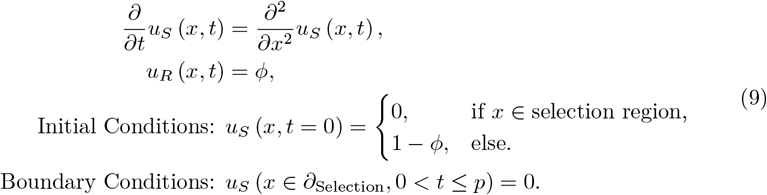

Over the time 0 *< t* ≤ *p*, the density profile of susceptible individuals smooths in the vicinity of the selection region, see Fig. 1a (*t* = *p*).

#### 2) Post-Selection Phase: *p < t <* 1

For the rest of the generation time, *p < t <* 1, the control measure is absent: the susceptible population is now capable of diffusing into the selection region without being removed. Given that once again the distribution of resistant individuals remains constant, the system now evolves according to:

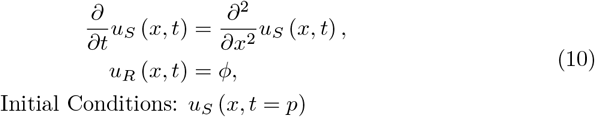

See Fig. 1a for an illustration of these distributions at *t* = 1.

#### 3) Breeding Phase: *t* = 1

At *t* = 1, the population undertakes a breeding step, in which the total population density is brought up to the carrying capacity of 1, whilst maintaining the local proportion or frequency of susceptible and resistance individuals. With susceptible and resistant population densities after breeding denoted as *U*_*S*_(*x*) and *U*_*R*_(*x*), respectively, the breeding step is formally defined as:

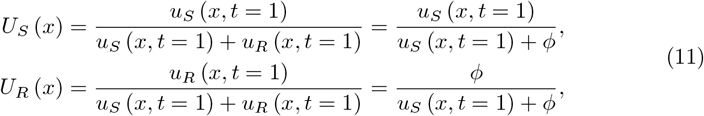

where we used the fact that *u*_*R*_(*x, t*) = *ϕ* for all *t* up to breeding. See Fig. 1a for illustrations of both *U*_*S*_ (*x*) and *U*_*R*_ (*x*).

#### Quantifying Rise in Resistance

The presence of a temporary, spatially isolated selection region prompts a local increase in the number of resistant individuals within and in the vicinity of the selection region (compare the initial and final resistant population densities in Fig. 1a). The change in the number of resistant individuals can be quantified by integrating the difference between the final and initial resistant population densities over the whole domain:

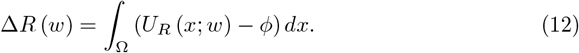

We numerically solve this integral with the SciPy [Vir+20] implementation of Simpson’s method.

### S2: Isolated

The control measure is applied in the selection region, 0 ≤ x ≤ *w*, while the refuge regions extend to infinity on both sides, i.e., there is refuge for *x <* 0 and *x > w*.

#### 1) Isolated Selection Phase: 0 *< t ≤ p*

The evolution of the system while the control measure is active is governed by the diffusion equation, with the following initial and boundary conditions:

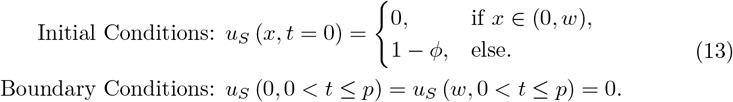

We obtain *u*_*S*_ (*x, t* = *p*) by the method of Laplace transforms and contour integration, as discussed in a similar example on page 747 of Riley, Hobson, and Bence [RHB06]:

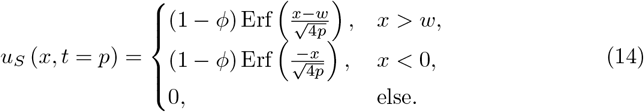

#### 2) Isolated Post-Selection Phase: *p < t <* 1

The evolution of the system during the remainder of the generation is governed by the diffusion equation subject to the initial condition:

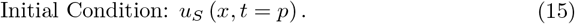

The solution at *t* = 1, *u*_*S*_ (*x, t* = 1), is given by the convolution of *u*_*S*_ (*x, t* = *p*) with the diffusion dispersal kernel:

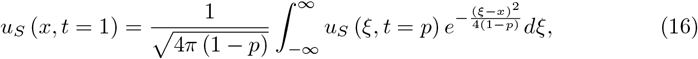

which is evaluated numerically by using the fast Fourier transform with NumPy [Har+20].

#### 3) Isolated Breeding Phase: *t* = 1

The post-breeding distribution of resistant individuals is given by Eqn. 11.

##### Quantifying Difference

From the post-breeding distribution, we define the total change in the number of resistant individuals for a given *w* as Eqn. 12:

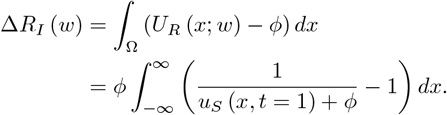

### S3: *w*_*U*_ Derivation

The solution to the susceptible distribution at the end of the post-selection phase for the isolated case in the limit of large *w* can be approximated by convolving the diffusion dispersal kernel with *u*_*S*_ (*x, t* = 1) when *w*≫ 1. The closed-form solution of this at the center of the selection region is given by:

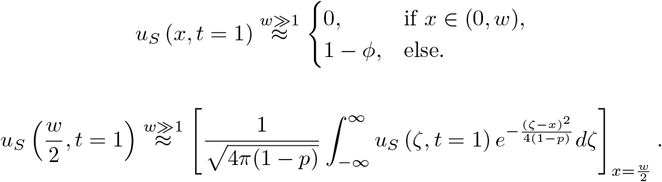

Due to symmetry, this can be solved by independently considering the evolution of the density distributions on the left (*x <* 0) and the right (*x > w*): The overall distribution will simply be the superposition of these two. Therefore, we define

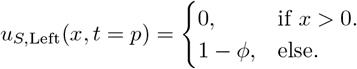

The density of susceptible individuals at *x* = *w/*2, *t* = 1 just from the left side is therefore:

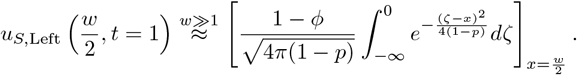

We employ the change in variable 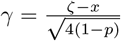, resulting in:

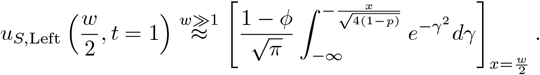

We can re-express the integral:

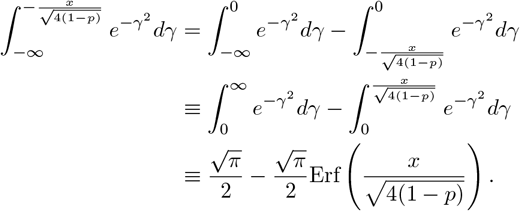

In which the second integral here comes from the definition of the error function [RHB06]. Taken together, we identify:

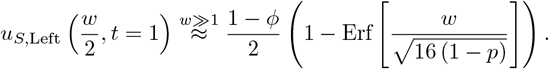

The contribution to the density of individuals at *x* = *w/*2, *t* = 1 from the right side (*x > w*) is the same as from the left. Therefore, the total contribution is:

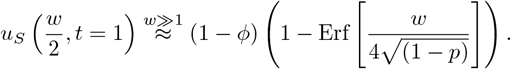

### S4: Periodic

#### 1) Periodic Selection Phase: 0 *< t ≤ p*

The evolution of the system while the control measure is active is governed by the diffusion equation, subject to the initial and boundary conditions:

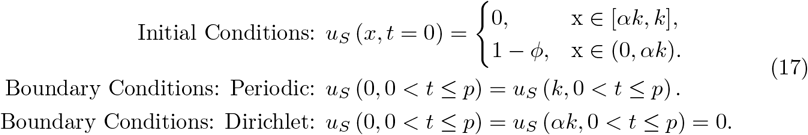

The solution of the diffusion equation, given the initial and boundary conditions at *t* = *p* can be obtained via the method of separation of variables (see Chapter 21 of Ref. [RHB06]):

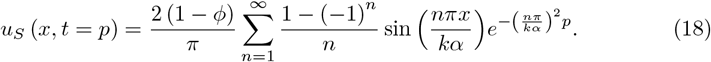

#### 2) Periodic Post-Selection Phase: *p < t <* 1

The evolution of the system during the remainder of the generation time is governed by the diffusion equation subject to the initial and boundary conditions:

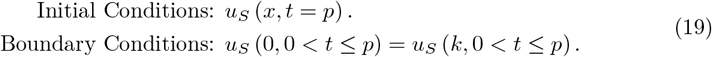

The solution to the diffusion equation, given the initial and boundary conditions, at *t* = 1 is obtained by the method of separation of variables (see Chapter 21 of Ref. [RHB06]):

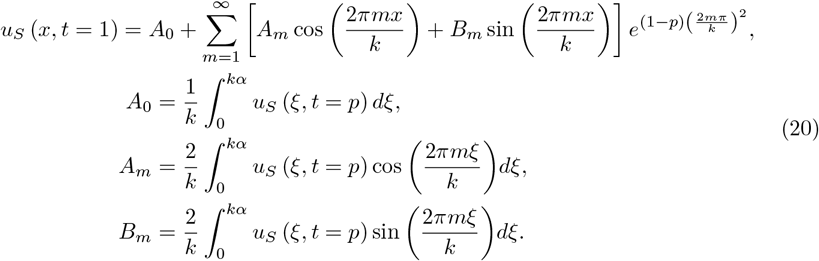

#### 3) Periodic Breeding Phase: *t* = 1

The post-breeding distribution of resistant individuals is given by Eqn. 11.

##### Quantifying Difference

From the post-breeding distribution, we define the total change in the resistant number over a single periodic sub-unit of width *k* as Eqn.12:

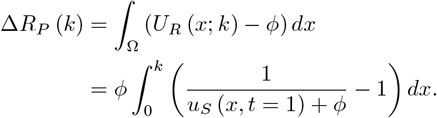

### S5: Multiple

Two possible multiple distributions are discussed in the main text: the unbound and bound scenarios. In both cases, the total amount of selection region, *c*, is divided into *N* regions of width *w*: we define the set of points which are in the selection region to be 𝒮 = {*x*| *x* = (*w* + *d*)*a* + *wb*}, such that *a* ∈ ℤ_*≥*0_ : *a < N* and *b* ∈ ℝ ∩ [0, 1]. In the case of the unbound scenario, the width of the refuge regions is constant, whereas for the bound case it varies with *N* (see Fig. 4). The following derivations are agnostic to the specific scenario considered.

#### 1) Multiple Selection Phase: 0 *< t ≤ p*

During this phase, the boundaries of the selection regions are Dirichlet boundary points: we shall formally define the set of these points to be 𝒟 = {*x*|*x* = (*w* + *d*)*a* + *wb*}, such that *a ∈ ℤ*_*≥*0_ : *a < N* and *b ∈* {0, 1}.

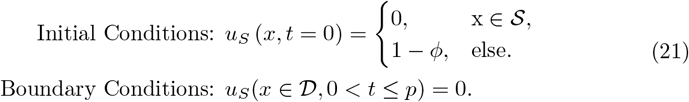

We shall define the set *d*_*M*_ = {*x*| (*x* mod (*w* + *d*) *> w* and 0 ≤ x ≤ Nw + (N − 1)d)} to be all the points of refuge sandwiched between selection regions, and define *E* = *Nw* + (*N* − 1)*d* to be the rightmost point of selection region. We note the result below is a combination of the results from the isolated (for *x > w* and *x <* 0) and periodic (*x ∈ d*_*M*_) cases:

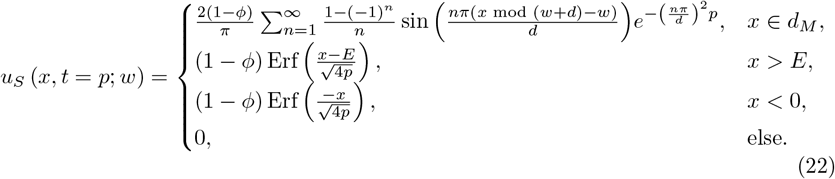

#### 2) Multiple Post-Selection Phase: *p < t <* 1

The evolution of the system during the remainder of the generation is governed by the diffusion equation subject to the initial condition:

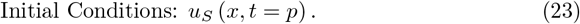

The solution to this at *t* = 1, is given by:

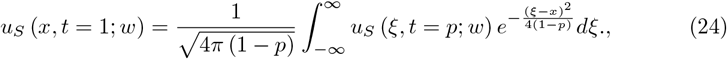

swhich is evaluated numerically by using the fast Fourier transform with NumPy [Har+20].

#### 3) Multiple Breeding Phase: *t* = 1

The post-breeding distribution of resistant individuals is given by Eqn. 11.

##### Quantifying Difference

From the post-breeding distribution, we define the total change in the resistant population as Eqn.12:

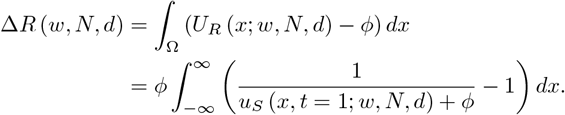

This is evaluated numerically with the SciPy [Vir+20] implementation of Simpson’s method.

